# Comparing the accuracy and efficiency of third generation DNA barcode sequencing: Oxford Nanopore Technologies versus Pacific Biosciences

**DOI:** 10.1101/2022.07.13.499863

**Authors:** Piotr Cuber, Darren Chooneea, Clementine Geeves, Silvia Salatino, Thomas J. Creedy, Claire Griffin, Laura Sivess, Ian Barnes, Ben Price, Raju Misra

**Affiliations:** Natural History Museum, London, SW7 5BD, UK; Concert Bio Translation and Innovation hub (I-HUB), Imperial College White City Campus, W12 0BZ, UK

## Abstract

At times of drastic decrease in biodiversity and loss of species, sometimes referred to as the “sixth mass extinction” or “Holocene extinction”, there is a high demand on the development of effective tools for studying and monitoring biodiversity. In the past decade, new promising technologies, such as third generation sequencing (TGS), enabled massive, rapid, and cost-effective data analysis of non-model organisms, accelerating taxonomic identification studies and contributing to conservation applications. Here, we focus on the comparison of the two main TGS providers, Pacific Biosciences (PacBio), and Oxford Nanopore Technologies (ONT), for the purpose of DNA barcoding. For ONT, we also tested selected combinations of different types of flow cells and ligation sequencing kits. Out of five tested combinations (PacBio, ONT Flongle flow cell & SQK-LSK110 kit, R9 flow cell & SQK-LSK109 kit, R9 & SQK-LSK100 kit, and R10 flow cell & Q20+ chemistry kit), ONT’s Flongle turned out to be most variable in returning the results, but at the same time the most cost efficient. The highest numbers of successfully sequenced samples were achieved with the ONT’s R10 & Q20+ chemistry combination. In terms of library preparation time, ONT protocols are the quickest, whereas regarding cost effectiveness - using Sanger pricing per sample as a cut-off - various technologies become affordable depending on the number of samples used. Although both tested platforms are suitable for DNA barcoding, we further discuss their limitations and applicability to different studies, with a special focus on the price and the number of samples. The pipeline we developed, from whole specimens to final DNA barcode consensuses, can aid planning and budgeting biodiversity studies, maximising the number of specimens sequenced in one run and speeding up the sample processing time.

## 1. INTRODUCTION

To date over 1.7 million species have been described by taxonomists around the world. However, this is just a small percentage of the millions of organisms on the planet [1]. The sheer scale of biodiversity is a great challenge to successfully classifying all species using traditional morphological based approaches. It has been estimated that the cost of describing five million animal species alone would stand at $250 billion and require six centuries to complete [2, 3].

Furthermore, morphological based taxonomy is fraught with limitations such as the difficulty delineating morphologically cryptic species and the shrinking number of talented taxonomists in the field. These limitations highlighted the need for more innovative approaches to taxonomy [4, 5]. In 2003 Paul Hebert et al. proposed a method of using genomic based approaches for taxonomic identifications [4]. This method was developed to provide rapid and accurate identification of organisms using short, conserved gene regions in various species, which was given the term DNA barcode [6]. Since then, DNA barcoding has become an established method for the rapid identification of biological organisms [3, 7].

Currently, first generation Sanger sequencing is the “golden standard” method for DNA barcoding projects. It has been found to remain a cost-effective method when sequencing fewer than 800 samples [8]. However, in the past few years, sequencing technologies have evolved rapidly. Ushering in high throughput, Next Generation Sequencing (NGS) technologies, such as Illumina or Ion Torrent, enabling users to multiplex hundreds of samples in a single sequencing experiment, driving down costs, while producing highly accurate sequencing data [9]. However, NGS sequencing technologies are limited by their ability to sequence short DNA fragments of around 100-600 bp which is smaller than most preferred DNA barcodes such as CO1 (650 bp) and therefore are suboptimal for accurate identification. More recently, high throughput third generation sequencing (TGS) platforms have been released, able to sequence longer reads, overcoming the limitations of NGS approaches while retaining their strength i.e., multiplexing hundreds of samples [10, 11]. Two third generation sequencers of note are the Sequel system by Pacific Biosciences (PacBio) and nanopore based sequencing developed by Oxford Nanopore Technologies (ONT).

The PacBio Sequel system (Sequel I, II, and IIe) use Single Molecule Real Time (SMRT) sequencing to generate highly accurate long read sequences. With SMRT sequencing, hairpin adaptors are ligated to double stranded DNA templates to form a closed, circularised template called SMRTbell libraries. Sequencing primers are annealed to the SMRTbell library to facilitate DNA polymerase binding, which acts as an anchor to the bottom of the sequencing cell and marks the initial site for sequencing. This circularised template is repeatedly sequenced to produce subreads. Consensus from these subreads generates a circular consensus sequencing read (CCS) which in turn produces a HiFi read with an accuracy of greater than 99.9% [7, 12, 13].

The ONT technology is based on nanopores, where the sample DNA is directly sequenced in real-time. It comes in various configurations of sequencing flow cells and chemistries. This includes scalable sequencers from the Flongle, a cheaper low-capacity alternative to the MinION flow cells. When sequencing, the strand of DNA from the sample is pulled into a biological nanopore and a minute electric current is generated. This electrical current is then converted using software to correspond to one of the bases on the sample DNA [14, 15]. The ONT now boasts a modal raw read accuracy of 99% using the new Q20+ chemistry and improved software.

## 2. THE AIM OF THE STUDY

The aim of this study was to compare the performance of TGS platforms, ONT and PacBio for barcode sequencing and evaluate their respective strengths and weaknesses for this purpose. We tested over 250 samples, performing technical replicates to determine reproducibility, from extracted DNA to library and technical library only replicates. We tested various iterations of flow cells available from the ONT at the time this paper was written (R9 and R10 MinION and Flongle) and sequencing kit chemistries (SQK-LSK109, SQK-LSK110, and Kit SQK-LSK112 early access Q20+) to see if they improved the demultiplexing and identification of these samples sequenced.

## 3. MATERIAL AND METHODS

### 3.1. Specimens’ description (collection, morphological analysis, plating)

Each specimen out of a total 262, was collected in the UK and identified by an expert taxonomist using standard methods and stored in 70-80 % ethanol. Before delivery to NHM some samples were stored at room temperature for an extended period, however on delivery to NHM all samples were stored at –20 °C. Each specimen was imaged before tissue sampling, typically a dorsal, ventral and/ or lateral view depending on the taxonomic group. Tissue was removed into the well of a PCR plate with 100 % ethanol (e.g., a leg from the insect specimens), unless the specimen was less than 5mm long in which case the whole body was used for DNA extraction. All equipment (forceps, petri dishes etc.) were cleaned with Trigene Advance disinfectant and rinsed in deionized water between specimens to prevent contamination, with forceps also heated in a bead heater for 10 - 15 seconds at 270 °C.

### 3.2. Molecular analysis

Molecular analysis workflow included several stages of laboratory and bioinformatic analyses developed in-house (Figure 1). The experiment was performed in two types of technical replicates, 3 repeats for each type, to identify potential weak points of the pipeline. One type of replicate involved partial pipeline repeats (PTR) starting with pooling stage to verify repeatability of next generation sequencing platforms (Figure 1), replicates were named Repeat1, Repeat2 and Repeat3. Another type called “full technical replicate” (FTR) started with the first PCR run and processed in 3 repeats (Figure 1), replicates were named PacBio1, PacBio2, PacBio3. Repeat1 was at the same time PacBio1.

**Figure 1.**
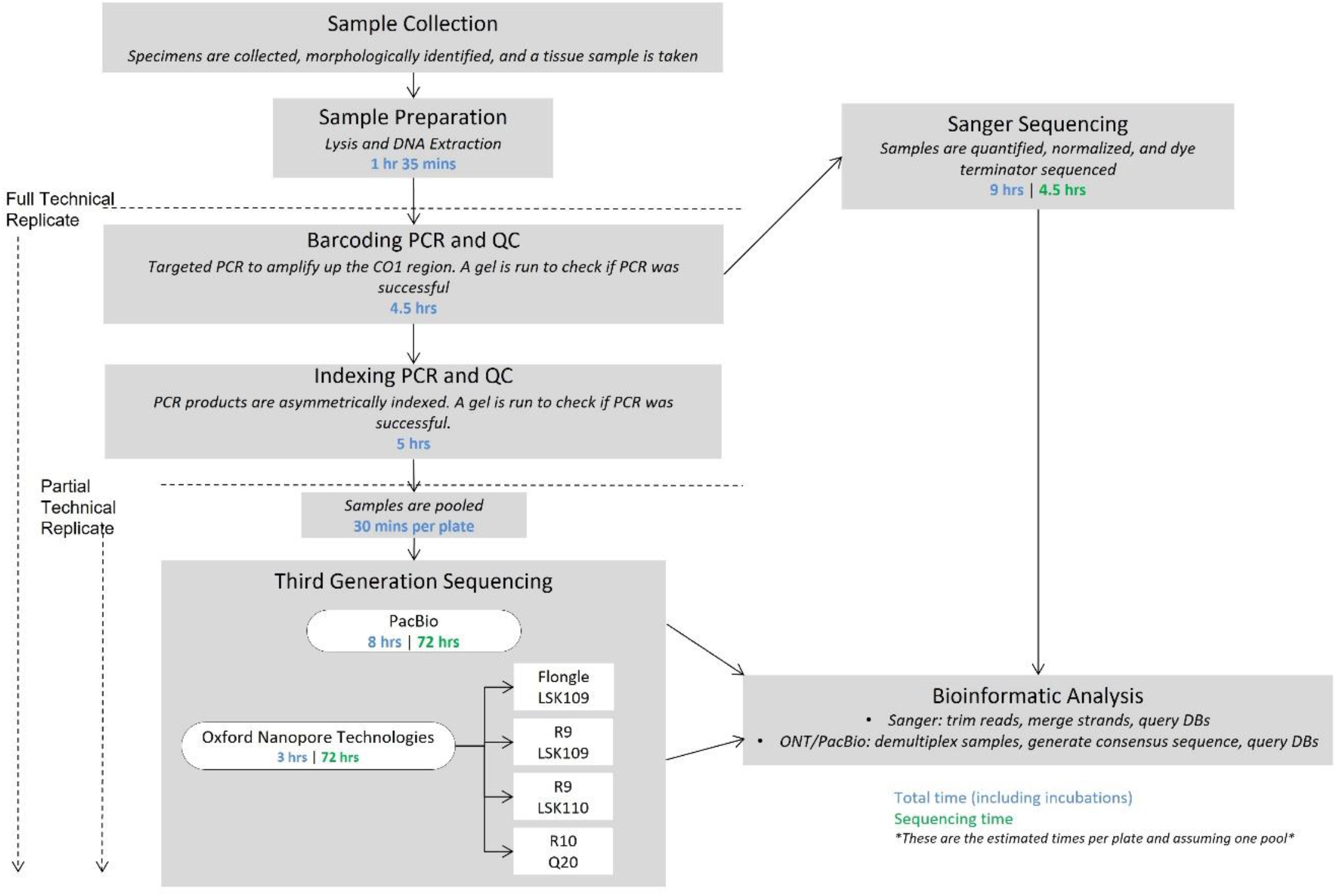
Experiment workflow. Diagrammatical representation of the comparison experiment. Timings are estimates of the total time at each step, including hands-on time and incubations, and are based off the assumption that one plate or pool is being processed at one time.

#### 3.2.1. DNA extraction, PCR, QC

The procedure began with the removal of as much ethanol as possible by pipetting without disrupting the specimen. Then the plate was placed on the thermocycler for 10 minutes at 70 °C (with the lids off and the thermocycler lid set to off) to evaporate any residual ethanol. The evaporation was prolonged if required. After this stage, the plate was removed from the thermocycler and the following was added; 5 µl 10X KAPA Express extraction buffer, 1 µl 1 U/µl KAPA Express extract enzyme, and 44 µl dH2O for a total volume of 50 µl per reaction. Specimens were pushed down with the end of the loop swab if necessary to ensure they were submerged by the extract mix, then the whole plate was spun down briefly. The DNA was extracted following the KAPA Express Extract kit protocol, with modifications in place to halve the reaction volume as well as in increased incubation time to allow the lysate mix to absorb into the soft tissue, and to avoid the use of proteinase K or manually squashing each individual specimen. The samples were incubated for 30 min at 30 °C, 60 °C for 30 min, 95 °C for 5 min and 4 °C hold. Post-lysis, the crude lysate was transferred to a new plate avoiding transferring any debris. To increase PCR success rate 1:5 lysate dilution can be used (unpublished data).

The KAPA2G Robust HotStart PCR Kit was used for PCR. The PCR reaction consisted of 5 µl Buffer B (5X), 5 µl Enhancer (5X), 0.5 µl 10 mM DNTPs, 1.25 µl 10 µM LepF Primer, 1.25 µl 10 µM LCO1490 Primer, 1.25 µl 10 µM LepR Primer, 1.25 µl 10 µM HCO2198 Primer, 0.1 µl DNA Polymerase (5 U/µl), and 4.4 µl PCR grade water per sample. 5 µl 1:5 dilution of the lysate was used to increase the success rate of PCR (unpublished data). The plate was placed in a thermocycler under the following conditions; initial incubation at 95 °C for 3 minutes followed by 5 cycles of 95 °C for 40 seconds, 45 °C for 40 seconds, 72 °C for 1 minute, then 35 cycles of 95 °C for 40 seconds, 55 °C for 40 seconds, 72 °C for 1 minute, followed by a final extension at 72 °C for 5 minutes and then a 4 °C hold.

The barcoding of metazoans makes use of the cytochrome c oxidase subunit 1 mitochondrial (CO1/COX1) gene. The proposed in-house developed universal primer cocktail consists of Folmer (LCO1490/HCO2198) and Lep (Lepidoptera) (Lep-F1/Lep-R1) primer sets (Table 1.). The sequences are shown in Table 1. Quality control check was performed by gel electrophoresis on 1% agarose gels run for 55-60 minutes. If any samples failed (i.e., no PCR band present) they were repeated with the same or a different primer set.

**Table 1.**
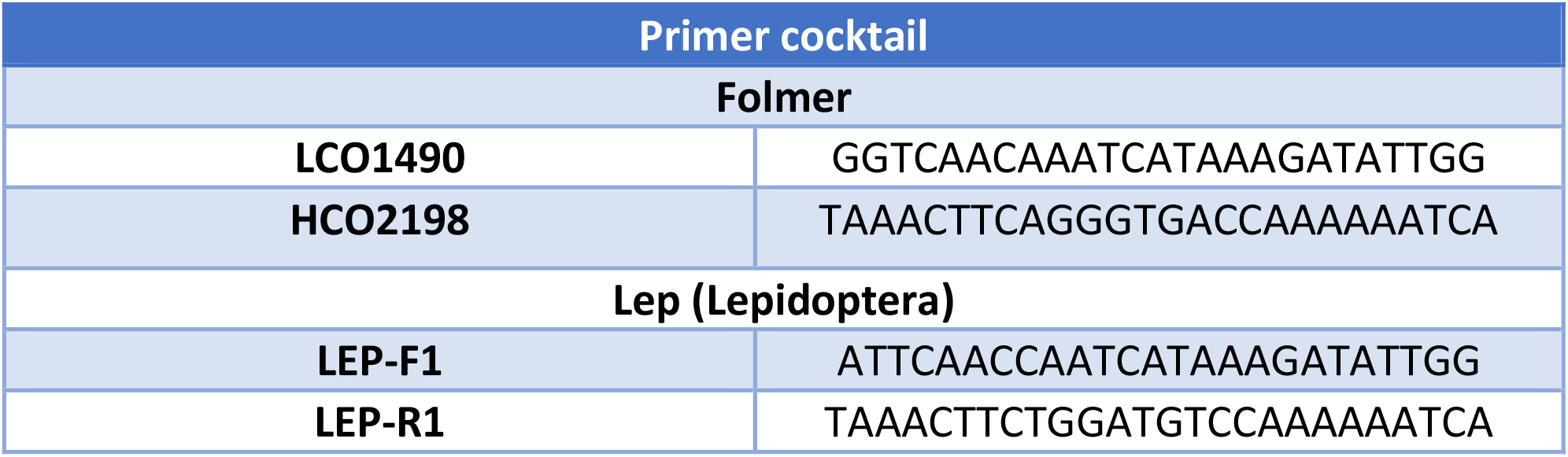
Primer cocktail used in the study [16, 17].

#### 3.2.2. Sanger sequencing

The Sanger Sequencing process makes multiple copies of a target DNA region, by amplifying products of various lengths, using the polymerase chain method. Running your PCR product out on an agarose gel checks for successful band amplification. In the second stage, the PCR product is combined in a tube or a plate, with the same primers used to generate the PCR product, dilution buffer, water, and BigDye Terminator V3.1 (this Kit contains DNA polymerase and four dye-labeled chain terminating dideoxy nucleotides). The DNA polymerase synthesizes new DNA, starting from the primer and continues to add nucleotide bases to the chain until a dideoxy nucleotide base is added. These chain terminating nucleotides (at the 3’ end) mark the end of each fragment and have a fluorescent dye incorporated, specific to that base. These extension products are then separated by capillary electrophoresis on a 3730xl DNA Analyzer. The fluorescently labeled chain terminating nucleotides excite the laser as they pass a tiny sensor allowing the end base to be identified and a DNA sequence to be generated.

PCR samples were first purified using AppMag magnetic PCR beads, following the manufacturer’s protocol. The concentration of each purified sample was then measured using a Nanodrop 8000 DNA spectrophotometer. Any samples outside the concentration required, for the fragment size amplified, were normalized. Forward and reverse samples were then set up for Sanger Sequencing using Applied Biosystems BigDye Terminator Kit V3.1 and cycled on a thermocycler, following an “in-house” adaptation of the manufacturer’s protocol using 28 cycles rather than the standard 25 cycles. Samples were then further purified to remove unincorporated dye terminators using AppMag Dye Terminator Removal Magnetic Beads, and an adapted method of the Axygen DyeClean protocol, using a volume of 5 μl instead of 10 μl per reaction. Purified reactions were loaded and sequenced on an Applied Biosystems 3730xl capillary DNA Analyzer. Sanger Sequencing can produce high quality sequence reads of up to 950 bp. The Folmer/Lep cocktail generates fragments around 750 bp.

#### 3.2.3. Sample preparation for third generation sequencing (TGS) platforms

The asymmetric indexing PCR of the barcoded products was carried out using the KAPA 2G Robust HotStart PCR Kit (KK5518). Indexes were ordered through IDT using PacBio’s multiplexing sequences (see appendix 1). Asymmetric indexing plates were created by using 24 Forward and 34 Reverse M13 tailed indexes (appendix 2). The following reagents were added to the indexing plates; 5 µl Buffer B (5X), 5 µl Enhancer (5X), 0.5 µl 10 nM dNTPs, 0.1 µl DNA Polymerase (5 U/µl), 8.9 µl Molecular Grade Water, and 3 µl template. The cycling conditions consisted of an initial incubation at 95 °C for 3 minutes followed by 5 cycles of 95 °C for 40 seconds, 45 °C for 40 seconds, 72 °C for 1 minute, then 35 cycles of 95 °C for 40 seconds, 55 °C for 40 seconds, 72 °C for 1 minute, followed by a final extension at 72 °C for 5 minutes and then a 4 °C hold.

Indexed PCR products were then analysed on a 1 % agarose gel. Samples that had a band present at the desired size (approximately 750 bp) were pooled by equal volumes (5 µl per sample). The pool cleaned was with AMPure® PB magnetic beads.

#### 3.2.4. Pacific Biosciences (PacBio) sequencing

SMRTbell libraries were produced following the “Procedure & Checklist - Preparing SMRTbell® Libraries using PacBio Barcoded M13 Primers for Multiplex SMRT® Sequencing” from section titled ‘Concentrate Pooled PCR Amplicons Using AMPure PB Beads’

The purified amplicons were diluted to a 500 ng input and underwent damage and end repair to form blunt ends to which hair pin adaptors were ligated to form the SMRTbell templates. The SMRTbell templates were purified with AMPure® PB magnetic beads after which the primer and polymerase were added to form the SMRTbell complex. The complex was loaded onto a single SMRT cell (SMRT Cell 1M) at 15pM for sequencing on a SEQUEL System using circular consensus sequencing (CCS) mode.

#### 3.2.5. Oxford Nanopore Technologies (ONT) sequencing

The ONT offers a range of combinations of flow cells with its sequencing kits and attached protocols. For the purpose of this experiment, we have chosen those most suitable in our judgement. We tested the following combinations:

- Flongle R9.4.1. (FLO-FLG001) called further simply Flongle used with Ligation Sequencing Kit (SQK-LSK110), called further, 110 and Amplicons by Ligation (SQK-LSK110) protocol version for Flongle.
- MinION R9.4.1. (FLO-MIN106D) called further R9 used with 110 kit and Amplicons by Ligation (SQK-LSK110) protocol.
- R9 used with Ligation Sequencing Kit (SQK-LSK109), called further 109, and Amplicons by Ligation (SQK-LSK109) protocol.
- MinION R10.4 (FLO-MIN112) called further R10 with Ligation Sequencing Kit (SQK-LSK112/Q20+), called further Q20 kit (SQK-LSK112) called further Q20 chemistry kit, and Amplicons by Ligation (SQK-LSK112) protocol.

SFB Expansion Kit (EXP-SFB001) and Flow cell Priming Kit (EXP-FLP002) were also used. The use of additional modules from New England Labs (NEB) is required by the attached protocols: NEBNext Companion Module for Oxford Nanopore Technologies Ligation Sequencing and NEB Blunt/TA Ligase Module. Sequencing reactions were performed on GridION Mk1device called further GridION, and MinKNOW software.

#### 3.2.6. Bioinformatic analysis

AB1 trace files of Sanger raw data were used as input for an in-house developed Nextflow [18] analysis pipeline to trim primers and poor base quality ends (with Trimmomatic version 0.39 [19], using the options “PE -phred33 HEADCROP:25 LEADING:20 TRAILING:20”), merge forward and reverse strands (with Pipebar version 3.00 [20], using the options “--mo 10 --ms 0.7”), and query the resulting sequences against the BOLD database (with bold_identification [21], using the options “-f fasta -d COX1 -n 20”) and against the non-redundant NCBI nucleotide collection database (with Blastn [22], using the options “-task megablast -db nt - remote -num_alignments 100”).

Long reads from Nanopore and PacBio sequencers were demultiplexed with ONTbarcoder [7] using default settings. The resulting FASTA files were queried against BOLD and NCBI databases with the same settings as for Sanger data. Finally, observed taxa were compared with the expected species, first using an in-house developed Python script that only kept the best matches in terms of “Similarity %” for BOLD and “Percentage of identical matches” for NCBI, and then doing a manual inspection to fix any misspellings, synonymous names, etc. We decided to follow a very strict approach regarding the manual inspection, qualifying the result as a “pass” only if it exactly matched the expected name after filtering out mentioned differences i.e., even if the genus name matched and either NCBI or BOLD databases records didn’t specify the name to the species level, it was classified as a “fail”. Private BOLD sequences were also manually classified as a “fail”, due to no means of verifying visually the full taxon name.

### 3.3. Phylogeny

To validate the accuracy and consistency of sequence recovery among sequencing methods and full/partial technical replicates, all unique sequences output from all methods were collated and dereplicated to find the exact sequence variants (ESVs), with the methodological conditions resulting in of each ESV recorded. The ESVs were then aligned using the MAFFT ginsi algorithm [23] based on a) the nucleotide sequences and b) the amino acid translation, after first assigning each sequence a reading frame and translating, then backtranslating after alignment, all using scripts from the biotools repository [24]. In R (R Core Team (2022). R: A language and environment for statistical computing. R Foundation for Statistical Computing, Vienna, Austria. URL, https://www.R-project.org/.) this alignment was then used to compute a distance matrix based on Fieselstein’s distance [25], which in turn was used to build a UPGMA phylogeny. These steps used the dist.dna and upgma functions from the ape [26] library, implemented in the maketree.R script of the metamate package (https://github.com/tjcreedy/metamate). The phylogeny was then processed using the recorded methodological conditions, first finding the most frequent ESV for a single DNA source well, then converting all other ESV terminals from that same source, representing sequences from different methods and replicates, into zero-height multifurcating subtrees to illustrate any divergence of sequences from different methodological conditions.

To further illustrate sequence divergence among methods and replicates, the distance matrix was ordinated using PCA, implemented in the princomp method in R. To control for differences among sequences from different species, the set of eigenvalues from different methods and replicates within each set of sequences from a source well were standardised to unit mean, then the first two PCA axes were plotted in a biplot.

## 4. RESULTS

### 4.1. Comparison of TGS technologies sequence generation capability

A total of 262 invertebrate specimens were selected for the comparisons. The morphological determination found the specimens belonged to 228 species, 145 genera, 73 families, 20 orders, 7 classes, and 3 phyla.

A total of 66 samples failed initial PCR stage (Figure 1.) for various replicates. 42 PCR reactions failed for FTR1 and all PTRs, 27 PCR reactions failed for FTR2 and 25 for FTR3. This stage was identified as the second most variable in the pipeline after demultiplexing stage. Different samples randomly failed demultiplexing for different replicates. The highest demultiplexing failure rate was found for Flongle (16.82% - 25.91%) and the lowest for R10 flow cell with Q20 chemistry (7.27% - 8.18%). This applied also to full replicates. Notably, Sanger produced the lowest failure rate (4.55%) in merging the sequences out of all technologies and throughout all the replicates.

It can be observed from Figure 2, for both types of replicates, that ONT’s Flongle turned out to be most variable in regard to number of samples successfully sequenced. The highest numbers of successfully sequenced samples were achieved with the ONT’s R10 & Q20+ chemistry combination followed by R9 combined with 2 types of kits and PacBio. Interestingly the difference in the proportion of successfully sequenced samples was only very slight in the case of R9 flow cell when using SQK-LSK109 kit and its updated version of SQK-LSK110 kit.

**Figure 2.**
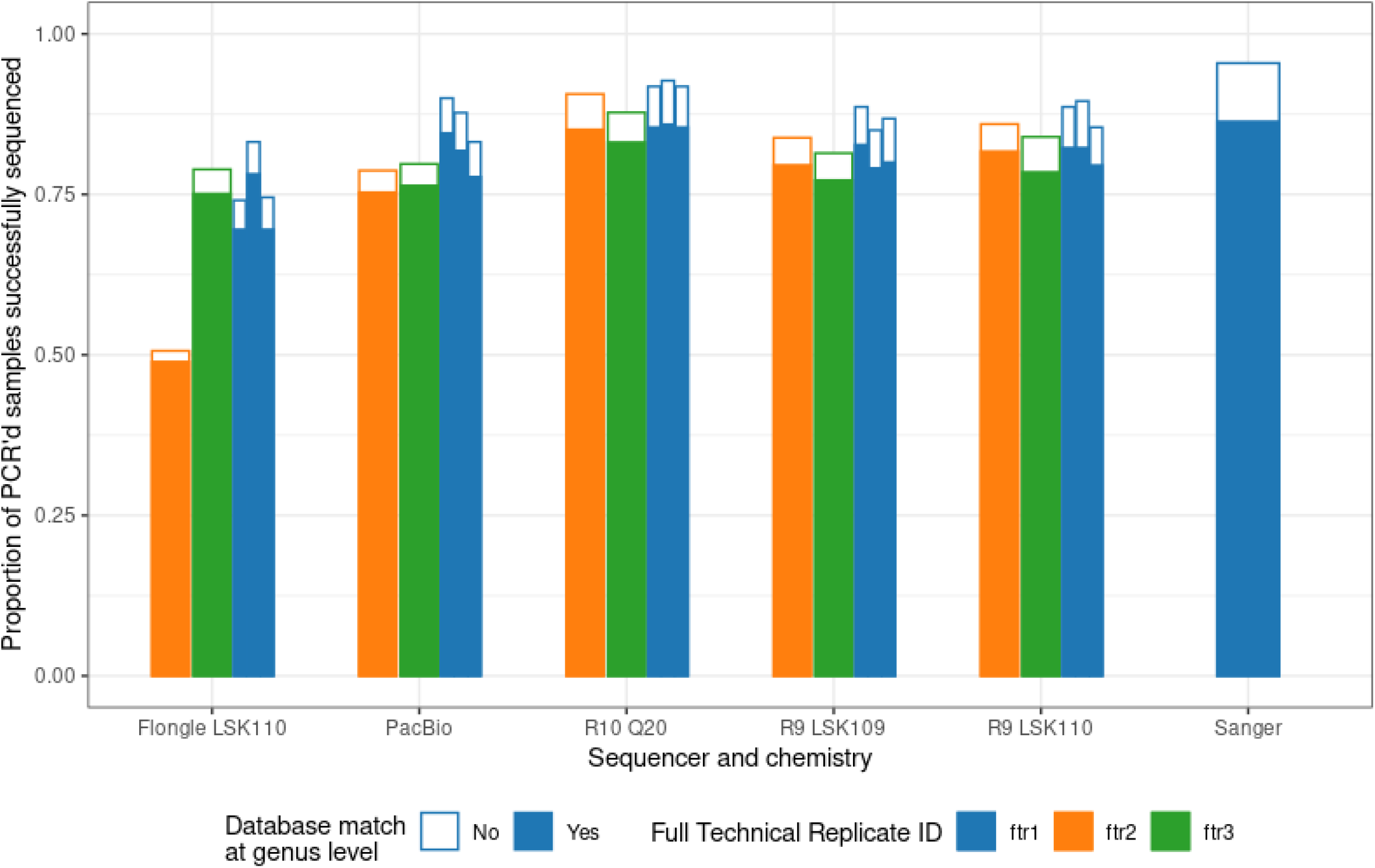
The proportion of full (FTR) and partial technical replicates (PTRs within FTR1) successfully returning a complete sequence for different sequencing methods, out of the 220, 235 and 237 successful PCRs (from 262 specimens total) in each separate FTR. Filled portion of bars represent the proportion of sequences that correctly matched against NCBI or BOLD at genus level. For TGS methods, the lack of a sequence for a given specimen after demultiplexing denoted failure; for Sanger, passing sequences were those that successfully merged the forward and reverse reads.

### 4.2. Phylogeny

From the 6143 sequences resulting from the different source DNA wells, methods, and replicates, 498 exact sequence variants (ESVs) were found. On inspecting the alignments, it was clear that the sequences from high throughput methods were very clean and consistent, but many of the Sanger sequences had single base insertions or deletions that resulted in frameshifts and erroneous amino acid alignments. Thus, the nucleotide alignment was used for phylogeny.

Figure 3 illustrates some typical and atypical observations from the phylogeny. Panels A and C show typical results; in many specimens, such as *Haliplus ruficollis* (panel A), *Leuctra inermis* and *Isoperla grammatica* (panel B), all high-throughput sequencing experiments resulted in the same sequence, while Sanger sequencing produced a similar sequence. In other specimens, such as *Gyrinus distinctus* (panel A) and *Leuctra hippopus* (panel C), there was more diversity among sequences from different experiments, although all sequences were more similar to one another than those from any other specimen. In a more atypical result, few specimens, such as *Agabus didymus* (panel A), reported that all 9 experiments using the DNA from this specimen generated the same sequence – however, in this case Sanger sequencing failed to generate a sequence. Other atypical results are illustrated in panels B and D of Figure 3. In in the top clade in panel B, the majority of ESVs are assigned to one species, *Baetis rhodani*, but are expected to be a range of different species from different taxa. In particular, note the ESVs of *Plectrocnemia conspersa* in the two clades of panel B; in 14 of the 19 experiments the resulting ESV was identical and was classified using BOLD as *Baetis rhodani* (top). But in the remaining 5 experiments, the resulting ESVs were classified as expected (bottom) and came out in the expected Trichoptera clade. Finally, clade D shows a number of specimens that returned sequences classified by BOLD as *Wolbachia* and other bacteria or symbiotic, commensal or parasitic organisms (e.g., *Chaetogaster limnaei* sequences were returned consistently for the samples identified as *Acroloxus lacustris*). In some cases, particularly *Ophion spp*., most or all recovered sequences were bacteria, while in others no high throughput sequences were recovered at all and the Sanger sequence was bacteria. In a curious case, *Calopteryx splendens*, the Sanger sequence was classified as Lestoidea while the two recovered high throughput sequences were classified as *Leuctra fusca*.

**Figure 3.**
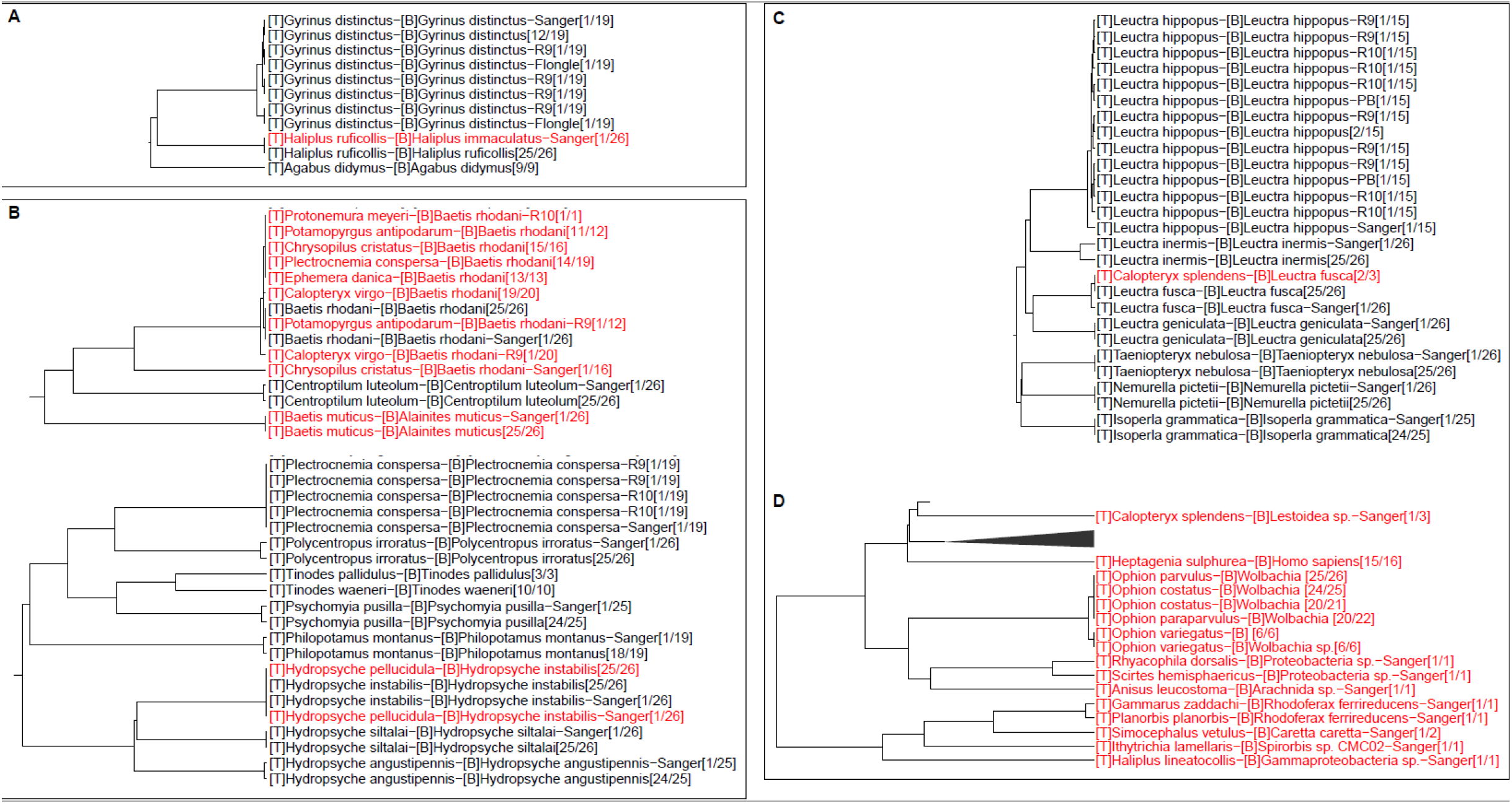
Selected clades from the UPGMA phylogeny of Exact Sequence Variants (ESVs) recovered from all methods (see supplementary Appendix 3 for complete tree). Tip labels show the expected species ([T}), species assigned by BOLD searches ([B]), the sequencing method (in some cases) and the number of different experiments that generated this ESV, as compared with the total number of experiments. Only sequencing methods that produced an ESV differing from the most common ESV are reported. For example, in panel A, 12 out of 19 experiments that used DNA from *Gyrinus distinctus* recovered the exact same ESV. The ESVs from the other 7 experiments are shown with the sequencing method reported. Tip labels coloured red denote a mismatch between expected and observed species identity. Panel A shows ESVs from three water beetle specimens (Coleoptera: Dytiscidae). Panel B shows two clades of ESVs that are classified using BOLD as Ephemeroptera (top) and Trichoptera (bottom). Panel C shows a clade of Plecoptera ESVs, and D assorted ESVs near to the root of the tree.

The ordination (Figure 4) shows that in the majority of cases, the sequences from different high throughput methods were very similar, with the majority of variation found between high throughput and Sanger sequences.

**Figure 4.**
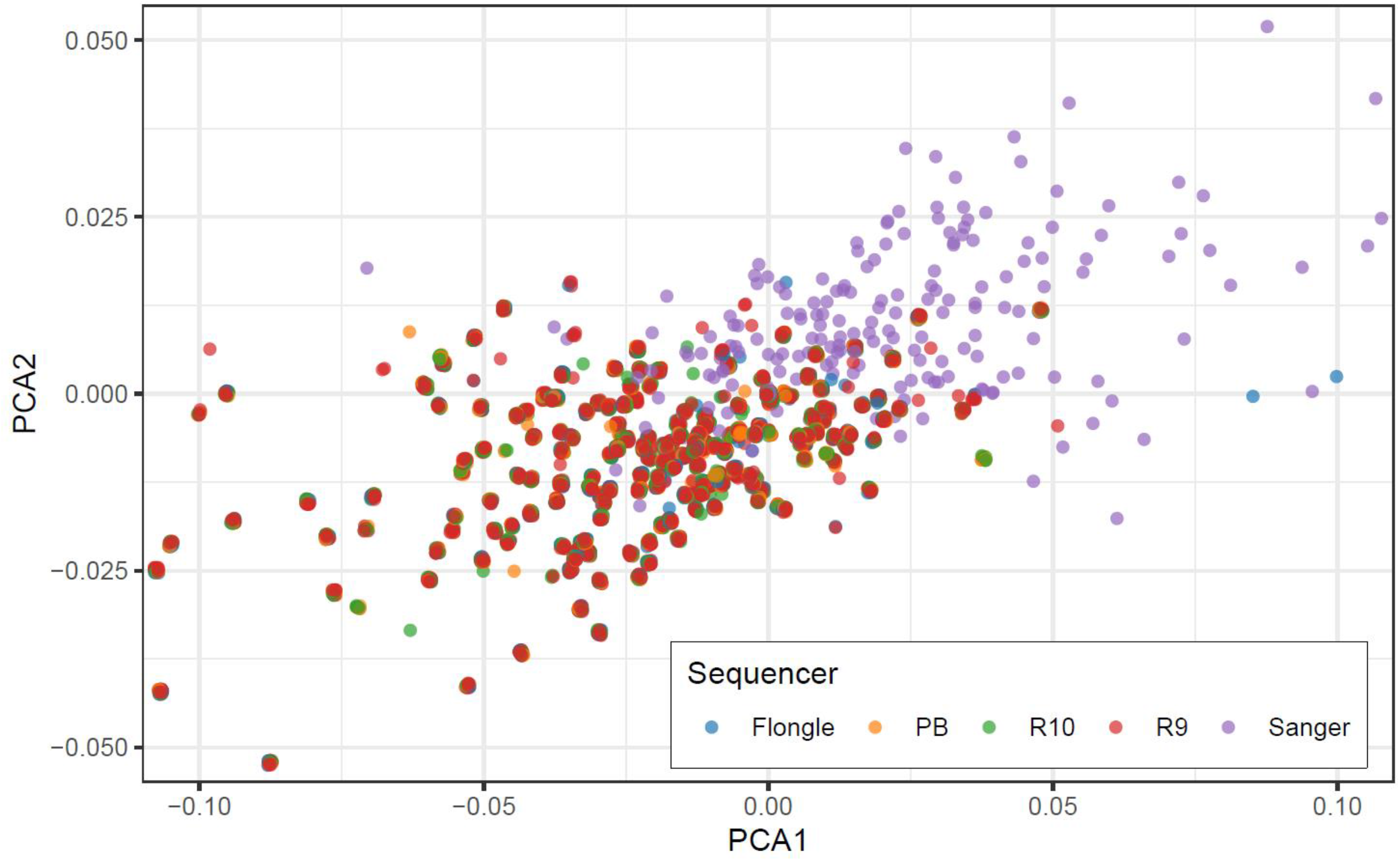
Ordination of pairwise distances among ESVs return from different experiments within source DNA. Colours denote sequencing method. Note that distances have been scaled to control for interspecific variation, and slight jitter has been applied to points to reduce overplotting. Several outliers are not shown.

### 4.3. Comparison of cost and time efficiency between TGS technologies

As seen on Figure 1 the most cost-effective protocols are those regarding the library preparation for the ONT platform (3 hours), whereas PacBio seems to be the most time-consuming procedure (8 hours) and one that is less intuitive and thus less user friendly. Interestingly both platforms have protocols quicker than sample preparation for Sanger (9 hours). Figure 5 presents cost-effectiveness of TGS technologies with Sanger arbitrary pricing of £6 per sample has been use as a cut off. ONT technologies become more affordable at lower number of samples in comparison to PacBio with ONT’s Flongle becoming most cost effective already at the number of 61 barcoded samples.

**Figure 5.**
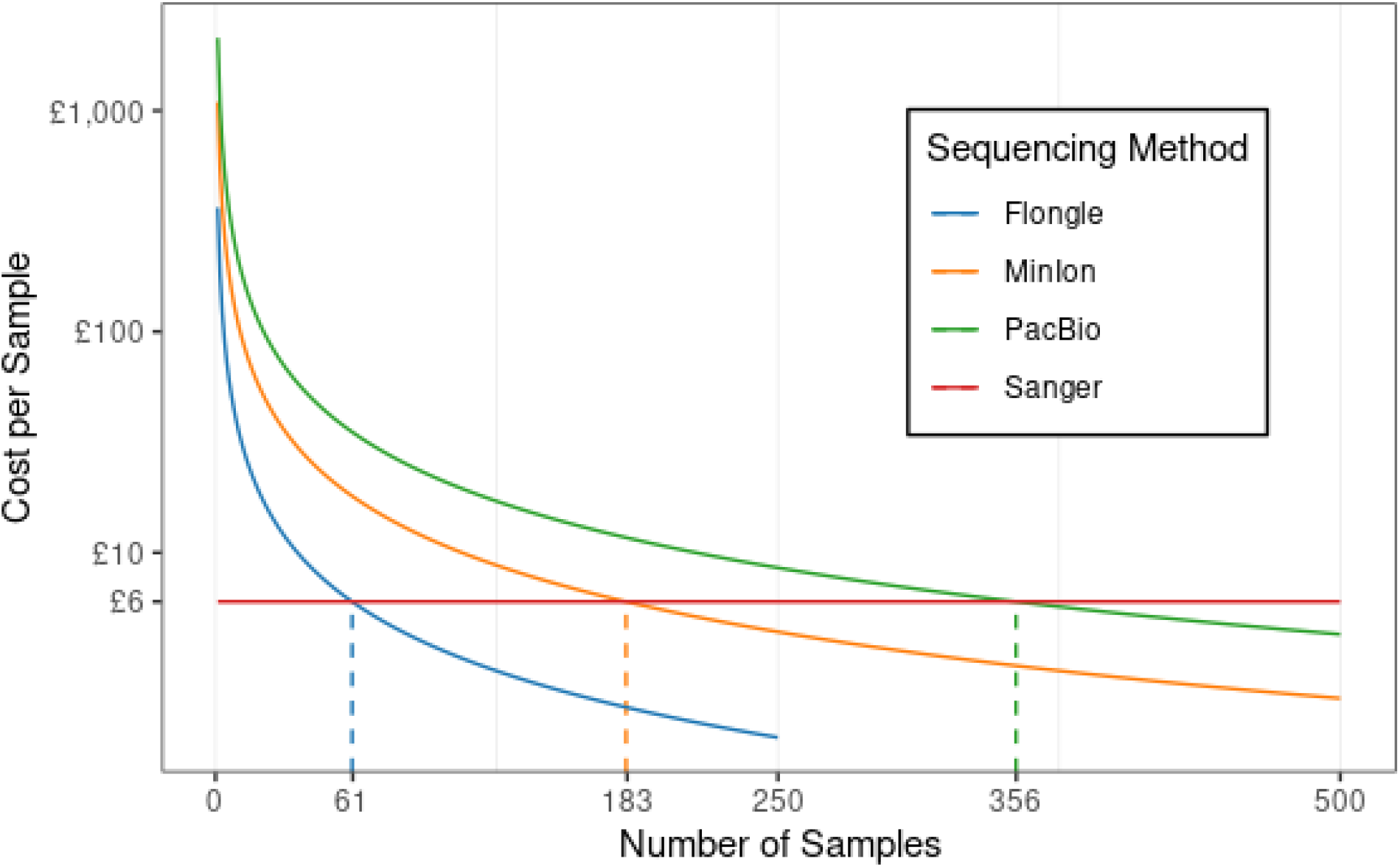
The cost per sample of different sequencing methods, based on the number of samples sharing a single sequencing run (i.e., flow cell or equivalent), up to the suggested maximum number of samples per flow cell (MinIon, PacBio not shown, both = 10,000). Dashed lines show the intersection of the cost/sample of TGS approaches with Sanger i.e., the break-even point above which it is cheaper per sample to sequence samples in parallel on a single flow cell on the given sequencer than to sequence each sample separately using Sanger.

## 5. DISCUSSION

Our study directly compares ONT with PacBio technologies for the application of barcode sequencing and species identification, building on the results obtained by Hebert at al. [8] regarding the PacBio and by Srivathsan et al. [7] for the ONT technologies individually. Here we directly compared the two technologies for the first time using morphological identification and Sanger sequencing as points of reference. Such comparison has only been described in depth between ONT and PacBio for metabarcoding purposes [10]. Although our study involved fewer specimens than two mentioned publications by Hebert et al. and Srivathsan et al. [7, 8], it includes greater diversity of species including insects’ and arachnids’ but also completely unrelated taxa such as leeches and gastropods. From the data presented here the best performing platform, when considering number of correctly determined identifications and cost was the ONT R10 flow cell using the Q20 chemistry kit, followed by the R9 flow cell (used with both kits), PacBio and Flongle. Once the library preparation process is considered, the ONT platforms were the best performing. As for the R9 flow cell, combination with SQK-LSK110 kit performed only marginally better than SQK-LSK109 kit confirming its improved accuracy.

TGS technologies become more economical with a higher number of samples/bigger projects, leveraging economies of scale. Although Sanger would remain the preferred method if the samples vs cost threshold for TGS technologies isn’t met. Here we found that threshold to be much lower than that calculated by Hebert at al. [8]. Out of the platforms tested, we found that Sanger is most cost effective when less than 60 samples are being analysed even when compared with Flongle, the smallest flow cell type available from ONT, which becomes cost-effective when 61 samples are being analysed. Other tested options only become competitive if more than 180 samples were being multiplexed (Figure 5). The same applies to time efficiency (Figure 1). Although the Flongle flow cell performed worse than other TGS technologies related to the proportion of successfully sequenced samples (Figure 2), it proved to be most cost-effective of the TSG options when investigating lower sample numbers i.e., between Sanger and the high-throughput technologies tested here. Several studies have suggested the upper limits for each technology type e.g., Srivathsan et al. suggested the upper limit for the Flongle at 250 samples [7]. MinION and PacBio on the other hand have the upper limit of 10,000 specimens [7, 8]. Together with the data presented here provide a guide to the optimal working range for each technology when considering cost vs number of samples.

If we compare turnaround time from DNA extraction to completed sequencing, when processing <4 plates, Sanger appears to be quicker, mainly due to the quicker sequencing time. >4 plates, both PacBio and ONT are far quicker due to the ability to multiplex samples together in a single sequencing run. If we were to use the one-step PCR method, where the barcodes are already indexed, it would reduce turnaround times further for each TGS platform to just 3 plates (96 samples) when compared to Sanger sequencing. When considering ease of use, Sanger being so well established is the most straight forward pipeline. But of the TGS platforms ONT is the easiest to generate a sequencing library from DNA extracts. PacBio is the more complicated of the three pipelines and has a longer library preparation time.

High quality barcode sequence data produced by TGS technologies has been well demonstrated [7, 8]. However, they primarily focused on insects and closely related taxa of arthropods, which are easier to process than many other types of invertebrates. Our study includes far more challenging taxa such as gastropods and are analysed here. The results of the phylogenetic and ordination examination of the sequence dissimilarities (Figures 3 and 4) showed that in most cases, there was either no or minimal variation among the different sequences found for a given specimen from TGS. In most cases, Sanger sequences diverged slightly; this is to be expected as the fragment length differed between TGS and Sanger sequencing. However, many Sanger sequences also contained small sequence errors. Atypical cases, where sequences from the same specimen diverged strongly, were shown through independent taxonomic classification of the sequences to be more likely due to cross-contamination of material (see the case of *Baetis rhodani*, Figure 3B), or the amplification of non-target DNA (Figure 3D), either from bacterial parasites or symbiotes, human contamination, or cross-contamination of the material during sampling prior to molecular investigation. Many of these cases only returned a sequence from Sanger sequencing, with TGS failing, or only a few TGS experiments succeeded: in these cases, it is likely that minimal target DNA was amplified, and the resulting Sanger sequence is a chimera of different non-targets (e.g., *Calopteryx splendens*, Figure 3C and D) and TGS sequencing and subsequent bioinformatic processing either failed to return an unequivocal read or returned an incorrect contaminant sequence.

The two PCR stages have been identified as areas that majorly affect the success rate for any sequencing platform used. Since this experiment, there have been a few ways in which to improve PCR success rate (unpublished data). Firstly, a 1:5 dilution of the lysate can help to dilute out contaminants and increase PCR yield. To go in hand with this, increasing the PCR cycle number to 40 cycles to compensate for the lower input. Secondly, using KAPA3G Plant PCR kit gives rise to a higher success rate. Thirdly, some specimens, notably gastropods or worms, are very viscous after extraction. This majorly impacts PCR therefore a 1X clean-up with Appmag or Seramag beads helps to decrease the fail rate for the viscous samples. Some samples become so viscous however that they prevent the migration of the magnetic beads and thus the clean-up. In these rare cases the clean up using Qiagen DNeasy PowerClean Pro Clean Up kit is recommended. As well as a clean-up on the gastropod samples, more specific primers can be used and to essentially keep repeating PCRs until they pass. Another improvement we found could make the pipeline even more time efficient is the use of primers that are already indexed and thus allow reducing the number of PCR reactions to only one. In terms of bioinformatic analysis, opportunities for future work include exploring other software, possibly all command-line based, to streamline the long-read data processing step and make it suitable also for high-performance cluster computing, thus saving time and computational resources.

In addition to the presented data, we evaluated other parameters. One initial experiment involved testing different library concentrations on the PacBio Sequel system. We tested three different loading concentrations, 12.5pM, 15pM, and 20pM, and found 15 pM gave the best initial results (unpublished data) and we decided to continue with this option. On the ONT platforms, we tested a combination of Flongle flow cell using SQK-LSK109 ligation kit, which resulted in failed QC (unpublished data) and consequently were not included in this analysis. We also tested the R10 flow cell with SQK-LSK110 kit, which gave comparable to R9 flow cell results (unpublished data), however, after releasing the Q20 chemistry kits this combination option is no longer supported by ONT and thus was excluded from the analysis too.

Both PacBio and ONT technologies are rapidly evolving, constantly improving and expanding what they have to offer. Therefore, some of the presented options, e.g., R9 flow cells from the ONT or Sequel system model from PacBio, might not be available in the future. But the data presented here, shows an optimistic direction of travel, where each technological advancement improves barcode sequencing applications. Furthermore, ONT have developed the PromethION family of sequencers with higher output flow cells, which support ultra-high throughput barcoding applications [15].

Finally, the presented studies confirm that curated databases such as BOLD ensure better results, compared to the larger NCBI database. However still require manual inspection of the results to correct errors such as spelling mistakes, present mostly in NCBI database but were not limited to it, e.g., *Netelia vibrator* instead of *Netelia virgata*. Although appears trivial, when automating pipelines and analysing greater number of samples impacts on misidentification rates. Also, often, if not for the manual curation of our reference dataset several identifications would have resolved only to the genus or higher taxonomic nomenclature. Here we were able to resolve to the species, and the results for species identification rates were similar between replicates but also between technologies. Another problem mentioned above is the use of taxonomic synonyms, in some cases it was more than two e.g., *Baetis* is synonymous with *Takobia* and *Alainites*. Interestingly, both databases were consistent with returning synonymous names e.g., BOLD database was returning name *Odeles marginata* whereas NCBI was returning *Elodes marginata*. Finally, there were also the cases where it was difficult to decide whether the author of the sequence misspelled the name, such as in the case of *Limnephilus luridus* and *Limnephilus lunatus*.

## 6. CONCLUSIONS

TGS platforms analysed in this study, both ONT and PacBio, are suitable for the purpose of barcode sequencing and thus for biodiversity research and monitoring. With sequencing costs lowering and newer technologies made available, sequencing accessibility is improving rapidly. Now, the biggest limitation for barcode sequencing applications is the cost and time taken for sample processing, whether it be manual or automated. Today, the quality of sequence data and the failure rate of producing sequences are no longer the limiting factors for any of the technologies tested. The pipelines and methods we developed here, from whole specimens to final DNA barcode consensuses, can aid planning and budgeting biodiversity studies when using either ONT or PacBio technologies, maximising their use for high throughput barcode sequencing of specimens in any laboratory.

## Ethical Approval

Not applicable

## Consent for publication

Not applicable

## Availability of data and metadata

BOLD and ENA databases, accession numbers pending.

## Author contribution

RM and DC designed the study, PC coordinated the study and manuscript writing, PC, CGe, DC and CGr performed laboratory side of the study, SS and TC performed bioinformatic side of the study. SS, TC, PC, DC and CGe performed result analysis and database preparation. TC and CGe prepared the figures. LS and BP plated the specimens and contributed to their morphological identification. All the authors contributed to writing and revising the manuscript.

## Acknowledgements

The authors would like to thank all the specialists who identified morphologically the specimens used in thus study.

## Appendixes

**Appendix 1.**
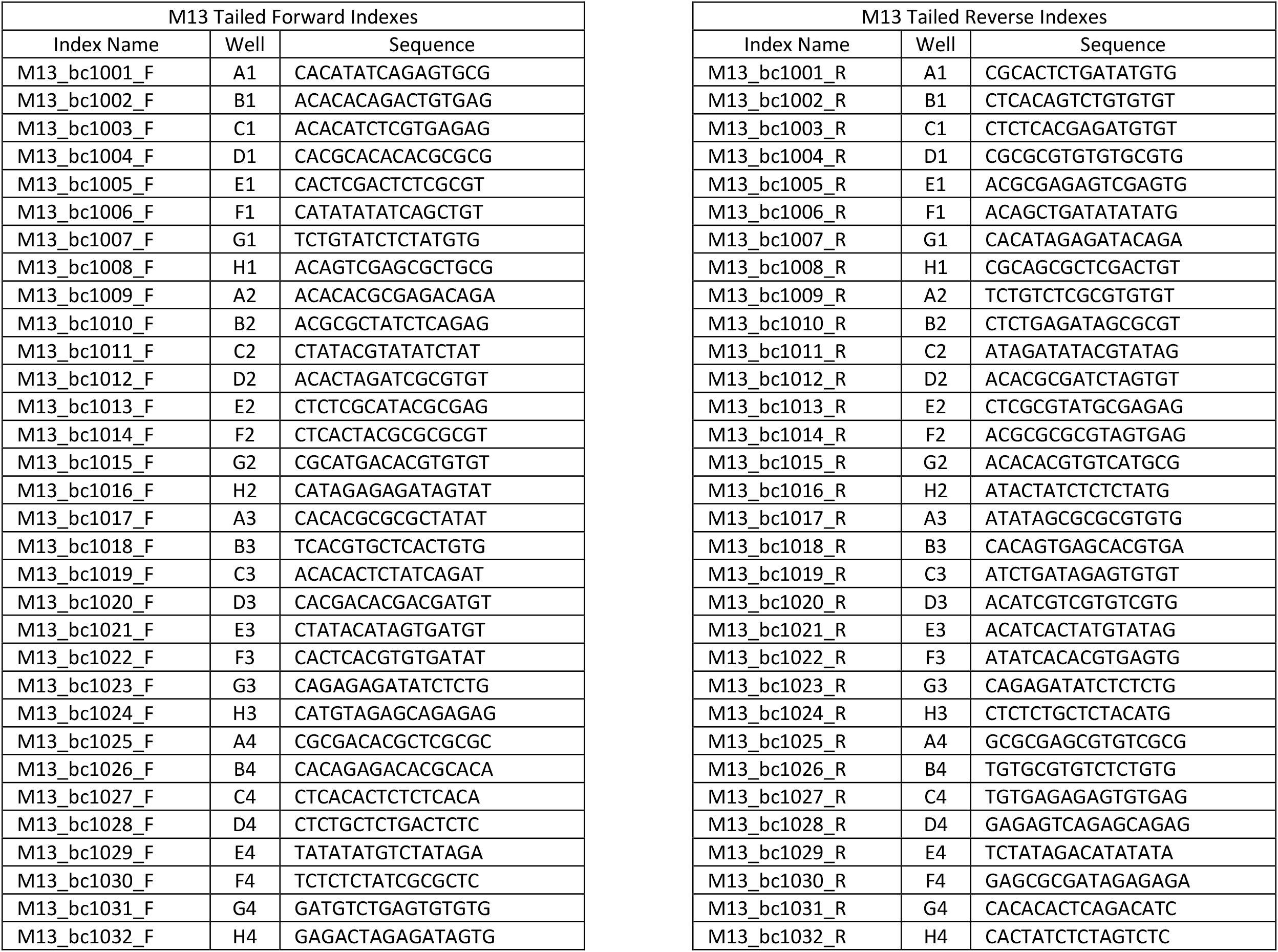
Indexing Sequences Below are the index sequences used. The indexes were ordered via IDT. The indexes were diluted to 10µM using dH2O.

**Appendix 2.** Asymmetric Indexing Matrix. 24 froward and 32 reverse indexes were used to create the indexing matrix. The matrix was designed to prevent reverse complementary pairings. 1.25µl of the 10µM dilution of a forward and a reverse index were combined in a well of a 96 well plate according to the tables below. The first well ID corresponds to the forward index and the second to the reverse e.g A1 – A2 is a combination of well A1 from the forward index plate and A2 from the reverse.

| Plate_1 |  |  |  |  |  |  |  |  |  |  |  |  |
| --- | --- | --- | --- | --- | --- | --- | --- | --- | --- | --- | --- | --- |
|  | 1 | 2 | 3 | 4 | 5 | 6 | 7 | 8 | 9 | 10 | 11 | 12 |
| A | A1-A2 | A1-B2 | A1-C2 | A1-D2 | A1-E2 | A1-F2 | A1-G2 | A1-H2 | A1-A3 | A1-B3 | A1-C3 | A1-D3 |
| B | B1-A2 | B1-B2 | B1-C2 | B1-D2 | B1-E2 | B1-F2 | B1-G2 | B1-H2 | B1-A3 | B1-B3 | B1-C3 | B1-D3 |
| C | C1-A2 | C1-B2 | C1-C2 | C1-D2 | C1-E2 | C1-F2 | C1-G2 | C1-H2 | C1-A3 | C1-B3 | C1-C3 | C1-D3 |
| D | D1-A2 | D1-B2 | D1-C2 | D1-D2 | D1-E2 | D1-F2 | D1-G2 | D1-H2 | D1-A3 | D1-B3 | D1-C3 | D1-D3 |
| E | E1-A2 | E1-B2 | E1-C2 | E1-D2 | E1-E2 | E1-F2 | E1-G2 | E1-H2 | E1-A3 | E1-B3 | E1-C3 | E1-D3 |
| **F** | F1-A2 | F1-B2 | F1-C2 | F1-D2 | F1-E2 | F1-F2 | F1-G2 | F1-H2 | F1-A3 | F1-B3 | F1-C3 | F1-D3 |
| **G** | G1-A2 | G1-B2 | G1-C2 | G1-D2 | G1-E2 | G1-F2 | G1-G2 | G1-H2 | G1-A3 | G1-B3 | G1-C3 | G1-D3 |
| **H** | H1-A2 | H1-B2 | H1-C2 | H1-D2 | H1-E2 | H1-F2 | H1-G2 | H1-H2 | H1-A3 | H1-B3 | H1-C3 | H1-D3 |

**Plate\_2**
|  | 1 | 2 | 3 | 4 | 5 | 6 | 7 | 8 | 9 | 10 | 11 | 12 |
| --- | --- | --- | --- | --- | --- | --- | --- | --- | --- | --- | --- | --- |
| **A** | A2-A4 | A2-B4 | A2-C4 | A2-D4 | A2-E4 | A2-F4 | A2-G4 | A2-H4 | A2-E3 | A2-F3 | A2-G3 | A2-H3 |
| **B** | B2-A4 | B2-B4 | B2-C4 | B2-D4 | B2-E4 | B2-F4 | B2-G4 | B2-H4 | B2-E3 | B2-F3 | B2-G3 | B2-H3 |
| **C** | C2-A4 | C2-B4 | C2-C4 | C2-D4 | C2-E4 | C2-F4 | C2-G4 | C2-H4 | C2-E3 | C2-F3 | C2-G3 | C2-H3 |
| **D** | D2-A4 | D2-B4 | D2-C4 | D2-D4 | D2-E4 | D2-F4 | D2-G4 | D2-H4 | D2-E3 | D2-F3 | D2-G3 | D2-H3 |
| **E** | E2-A4 | E2-B4 | E2-C4 | E2-D4 | E2-E4 | E2-F4 | E2-G4 | E2-H4 | E2-E3 | E2-F3 | E2-G3 | E2-H3 |
| **F** | F2-A4 | F2-B4 | F2-C4 | F2-D4 | F2-E4 | F2-F4 | F2-G4 | F2-H4 | F2-E3 | F2-F3 | F2-G3 | F2-H3 |
| **G** | G2-A4 | G2-B4 | G2-C4 | G2-D4 | G2-E4 | G2-F4 | G2-G4 | G2-H4 | G2-E3 | G2-F3 | G2-G3 | G2-H3 |
| **H** | H2-A4 | H2-B4 | H2-C4 | H2-D4 | H2-E4 | H2-F4 | H2-G4 | H2-H4 | H2-E3 | H2-F3 | H2-G3 | H2-H3 |

**Plate\_3**
|  | 1 | 2 | 3 | 4 | 5 | 6 | 7 | 8 | 9 | 10 | 11 | 12 |
| --- | --- | --- | --- | --- | --- | --- | --- | --- | --- | --- | --- | --- |
| **A** | A3-A1 | A3-B1 | A3-C1 | A3-D1 | A3-E1 | A3-F1 | A3-G1 | A3-H1 | A3-E4 | A3-F4 | A3-G4 | A3-H4 |
| **B** | B3-A1 | B3-B1 | B3-C1 | B3-D1 | B3-E1 | B3-F1 | B3-G1 | B3-H1 | B3-E4 | B3-F4 | B3-G4 | B3-H4 |
| **C** | C3-A1 | C3-B1 | C3-C1 | C3-D1 | C3-E1 | C3-F1 | C3-G1 | C3-H1 | C3-E4 | C3-F4 | C3-G4 | C3-H4 |
| **D** | D3-A1 | D3-B1 | D3-C1 | D3-D1 | D3-E1 | D3-F1 | D3-G1 | D3-H1 | D3-E4 | D3-F4 | D3-G4 | D3-H4 |
| **E** | E3-A1 | E3-B1 | E3-C1 | E3-D1 | E3-E1 | E3-F1 | E3-G1 | E3-H1 | E3-E4 | E3-F4 | E3-G4 | E3-H4 |
| **F** | F3-A1 | F3-B1 | F3-C1 | F3-D1 | F3-E1 | F3-F1 | F3-G1 | F3-H1 | F3-E4 | F3-F4 | F3-G4 | F3-H4 |
| **G** | G3-A1 | G3-B1 | G3-C1 | G3-D1 | G3-E1 | G3-F1 | G3-G1 | G3-H1 | G3-E4 | G3-F4 | G3-G4 | G3-H4 |
| **H** | H3-A1 | H3-B1 | H3-C1 | H3-D1 | H3-E1 | H3-F1 | H3-G1 | H3-H1 | H3-E4 | H3-F4 | H3-G4 | H3-H4 |

**Appendix 3.**
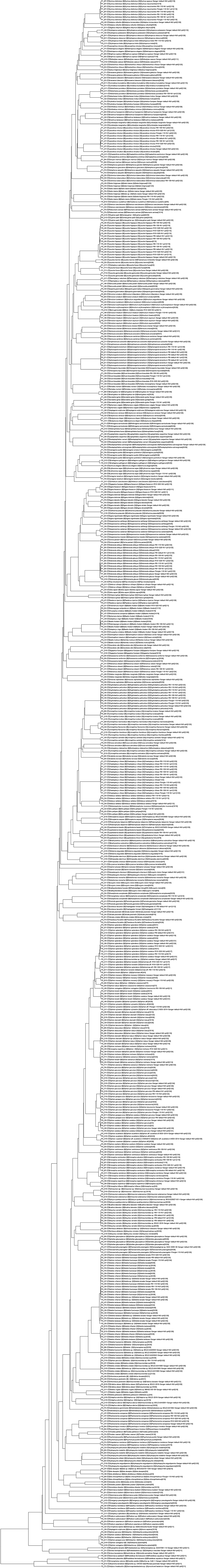
Complete UPGMA phylogeny of Exact Sequence Variants (ESVs) recovered from all methods

